# FSC-Q: A CryoEM map-to-atomic model quality validation based on the local Fourier Shell Correlation

**DOI:** 10.1101/2020.05.12.069831

**Authors:** Erney Ramírez-Aportela, David Maluenda, Yunior C. Fonseca, Pablo Conesa, Roberto Marabini, J. Bernard Heymann, Jose Maria Carazo, Carlos Oscar S. Sorzano

## Abstract

In recent years, advances in cryoEM have dramatically increased the resolution of Coulomb potential maps and, with it, the number of solved atomic models. It is widely accepted that the quality of cryoEM maps varies locally; therefore, the evaluation of the maps-derived structural models must be done locally as well. In this article, a method for the local analysis of the map-to-model fit is presented. The algorithm uses a comparison of two local resolution maps. The first is the local FSC (Fourier shell correlation) between the full map and the model, while the second is calculated between the half maps normally used in typical single particle analysis workflows. We call the new quality measure “FSC-Q”, and it is a quantitative estimation of how much of the model is supported by the signal content of the map. Furthermore, we show that FSC-Q may be helpful to avoid overfitting. It can be used to complement other methods, such as the Q-score method that estimates the resolvability of atoms.

## 1- Introduction

Single-particle cryo-electron microscopy (cryoEM) has become a powerful technique for the structural determination of biological macromolecules. In recent years, new direct detection cameras and better reconstruction algorithms have allowed to obtain Coulomb potential maps with a high level of detail. The next logical step is to build an atomic model that fits into the density map. Both, the quality of the reconstructed map and the derived model must be evaluated carefully to identify errors and poorly resolved regions. The quality of the maps is usually expressed in terms of a resolution measure. The typical resolution measure reported is based on a threshold in the Fourier shell correlation (FSC) curve between two independent reconstructions (Harauz and van Heel, 1986). The agreement of a model with a map can be done in a similar way by calculating the FSC between the experimental map and a map calculated from the model.

However, it is well known that the quality of a reconstruction depends on the region of the macromolecule and the resolution must be calculated locally (Cardone et al., 2013, Kucukelbir et al., 2014, Vilas et al., 2018, Ramirez-Aportela et al., 2019), and even directionally (Tan et al., 2017, Vilas et al., 2020). To evaluate the quality of an atomic model, different steric measurements have been integrated into Molprobity (Chen et al., 2010). However, these measures are based on the model itself, regardless of the cryoEM map. Local map-to-model fit measurements have been introduced, such as EMRinger (Barad et al., 2015), which takes into account density values near carbon-ß atom, and the recent Q-score (Pintilie et al., 2020), which measures the correlation between the map values at points around the atom and a reference Gaussian-like function.

In this paper we present a new measure (FSC-Q) for local quality estimation of the fit of the atomic model to the Coulomb potential map. The key difference with previously proposed methods is that we objectively and quantitatively estimate the specific parts of the model that are truly supported by the map on the bases of its local resolution. The rational of the method is based on the differences between local resolution values calculated with blocres (Cardone et al., 2013), considering the appropriate statistics to make each of the comparisons meaningful. The process involves the subtraction of the local resolution map between the final full map and the map generated from the atomic model, from the local resolution map between the two half-maps (where a half-map refers to a map reconstructed from half of the data set). This latter subtraction, when properly statistically scaled, provides an estimation of the signal content of the map itself, that is then compared with the signal content implied in the fitting of the structural model. When the fitting would require a signal content that, simply, is not present in the experimental data, FSC-Q flags it, providing in this way very objective information on how the structural model is supported, or not, by the local resolvability (the local resolution) present in the experimental map. It reveals potential modelling errors as well as poorly defined parts of models. The approach is conceptually simple, fully objective and quantitative, and lends itself very well to decisionmaking in a meta (or multi) criteria approach (Sorzano et al., 2020).

## 2- Methods

### 2.1. Local quality of fit

A common practice to globally assess the quality of the fit between the reconstructed map and the refined model is the use of the FSC. However, this practice only allows the fitting to be assessed as a whole. Here, we implemented an algorithm to evaluate the quality of the fitting of the atomic model into the density map following a similar strategy, but locally.

Our algorithm follows the general principle of making a comparison between the local FSC map calculated using the map obtained from the full data set and the map generated from the atomic model (***FSC_map–model_***). However, the key novelty is that this comparison is not performed involving directly the full reconstruction but, instead, it involves the local FSC map obtained when using the half maps (***FSC_half_***). We define FSC-Q as the difference between ***FSC_map–model_*** and ***FSC_half_***.

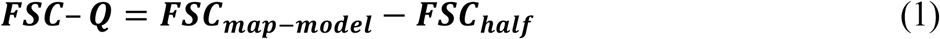

Obviously, this combination of local FSC’s calculated from half the data set and from the full data set requires the appropriate statistical analysis, that will be addressed in the next section. For the calculation indicated in Equation (1), a sliding window of size between five to seven times the overall resolution reported is used. A mask is applied to enclose the region of interest of the macromolecule. Those points of the map where there are “significant” differences between ***FSC_map–model_*** and ***FSC_half_***, allow us to detect errors in the refinement of the model or uncertainties in positions of parts of the model.

### 2.2. FSC threshold

Equation 1 compares an FSC determined from half the data set with an FSC obtained from the complete data set. Certainly this is not a simple issue and in the following we present how we perform this task using a rather simple and, we think, quite uncontroversial and widely accepted reasoning. We recall that (Cardone et al., 2013), proposed a FSC threshold of 0.5 to report the local resolution. We will start by choosing this threshold for the analysis of the FSC between two half maps. The approximate relationship between the FSC and the signal-to-noise ratio (SNR) in each of the half-maps is:

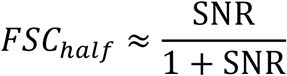

The threshold of 0.5 then corresponds to an SNR of 1. Note that this relationship is only approximate (Sorzano et al., 2017), but it can be taken as guidance to choose a FSC threshold related in some way to the corresponding SNR level. We may expand the relationship between the FSC and SNR to get

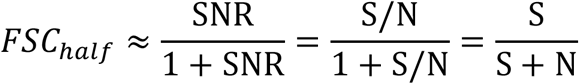

When we compare the full map versus the PDB converted to a map, the full map has half of the noise power than the two half maps, while the PDB does not have any noise. We may now reason

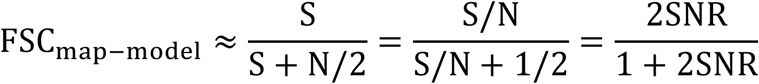

Choosing the same SNR value as for the half maps (SNR=1), we get a FSC threshold of ⅔. We must always bear in mind that this relationship is only valid on average and based on a 0-th order Taylor expansion (Sorzano et al., 2017), consequently its absolute values should not be taken too precisely. Indeed, any pair of thresholds can be used, as long as they represent a consistent SNR. Note that the distribution of FSC-Q values fluctuates around 0 when the map and model match (Supplementary Figure 1).

### 2.3. Input and output

The algorithm requires as input a 3D cryoEM density map, the fitted atomic model, the two half maps, and a mask enclosing the macromolecule. The blocres program (Cardone et al., 2013) is used to calculate the *FSC_map–model_* and *FSC_half_*. To assess the quality of the model fit, the *FSC_half_* map is subtracted from the *FSC_map–model_* map (equation 1), generating a difference map. However, we have found it useful to project this difference map onto atomic models (PDB format). We created a specific tool xmipp_pdb_label_from_volume, in XMIPP3 (de la Rosa-Trevin et al., 2013); which represents the map values in the occupancy column for each atom in the model.

In all the cases analyzed in this work, the function xmipp_volume_from_pdb (Sorzano et al., 2015) was used to generate the density map from the atomic model. From this map, a mask was generated using the xmipp–create 3d mask protocol in *Scipion* (de la Rosa-Trevin et al., 2016). A threshold of 0.02 and a dilation of 3 pixels were applied. In all cases, a sliding window of five times the reported resolution was used.

### 2.4. Code availability

The protocol used here has been integrated into the image-processing framework *Scipion* (de la Rosa-Trevin et al., 2016), in the development branch https://github.com/I2PC/scipion-em-xmipp (this branch will eventually become the next release of *Scipion3*). Different visualization options have been implemented to analyze the results. For example, the display of the map or the atomic model in UCSF Chimera (Pettersen et al., 2004) colored according to the calculated FSC-Q values.

## 3- Results

### 3.1. Fitting Analysis

To test our algorithm, we initially used the known 20S proteasome structure with a global resolution of 2.8 Å as reported by the gold-standard FSC of 0.143 (emd-6287) (Campbell et al., 2015). Figure 1 shows the FSC-Q values represented on a map generated from the atomic model (Figure 1A) or on the atomic model itself (PDB: id-6bdf) (Figure 1B). Atoms with FSC-Q close to zero means that the model is supported by the signal in the half-maps, while positive values far from zero corresponds to parts of the model that have a poor fit or that corresponds to areas in which the map had a poorer resolvability. Negative values correspond to areas where atoms are correlating with noise, or where, in general, there is not enough information to support the model. In this specific case there are few negative values and they correspond to external atoms of some side chains where there is no signal in the map.

**Figure 1.**
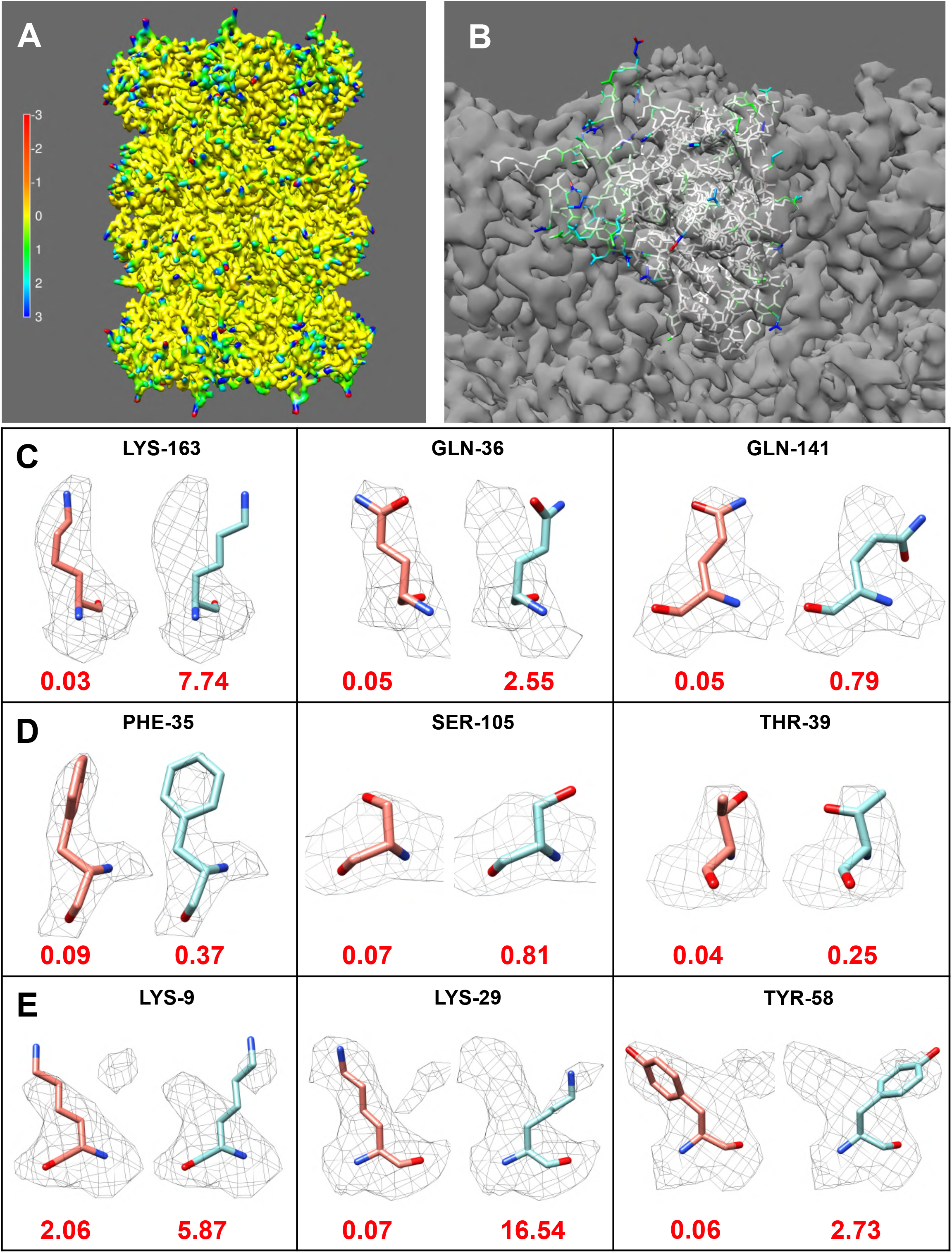
Detection of map-model fit errors using FSC-Q. FSC-Q values calculated for the 20S proteasome structure (emd-6287) are represented on the map generated from the atomic model (pdb id: 6bdf) (A) or on the atoms of chain Q of the atomic model (B). Panels C, D and E show rotamers of several amino acids in the 20S proteasome structure that have been altered. In each panel, the original rotamer is shown on the left and the modified rotamer on the right with their corresponding average FSC-Q scores. For the calculation of the average, the absolute FSC-Q value of each atom was considered. C) Rotamers of 3 long-chain amino acids that are clearly out of the density [LYS-163 chain B, GLN-36 chain V and GLN-141 chain T]. D) Very subtle modification of the residues: PHE-35 chain 1, SER-105 chain Q and THR-39 chain R. E) Residues in which the modified rotamers overlap with other densities [LYS-9 chain L, LYS-29 chain R and THR-58 chain Z].

To test the usefulness of this measure, we artificially modified the side chain rotamers of some amino acids of the 20S proteasome using Coot (Emsley et al., 2010). Figure 1C, D and E shows the altered amino acids divided into 3 blocks. A first block (Figure 1C) is made up of 3 long-chain amino acids (LYS and GLN) that are clearly out of the density. In the second block (Figure 1D) there are very subtle modifications and short side chains errors such as SER and GLN; finally, in the third block (Figure 1E) we show cases in which the modified rotamers overlap with other densities. In all cases the FSC-Q of the side chain increased compared to the original rotamer. This shows that even subtle changes such as aromatic ring rotation of the PHE-35 or side chain rotation of the SER-105 can indeed be detected by our method.

Furthermore, we analyzed the recent structure of the spike glycoprotein of SARS-CoV-2 in the closed state (emd-21452), using half-maps kindly provided by Prof. David Veesler (Walls et al., 2020). Two refined models were obtained from the repository of Prof. Andrea Thorn at https://github.com/thorn-lab/coronavirus_structural_task_force, the one proposed by the authors and a model corrected using ISOLDE (Croll, 2018). As expected, the corrected model as a whole has a lower average FSC-Q (FSC-Q = 0.96) compared to the original model (FSC-Q = 0.99). Furthermore, our method identified some regions, such as amino acids LEU-533 to VAL-534 and THR-323 to GLU-324, where the corrected model clearly fits better to the experimental data than the original model (Figure 2). In both areas a considerable reduction of FSC-Q is observed.

**Figure 2.**
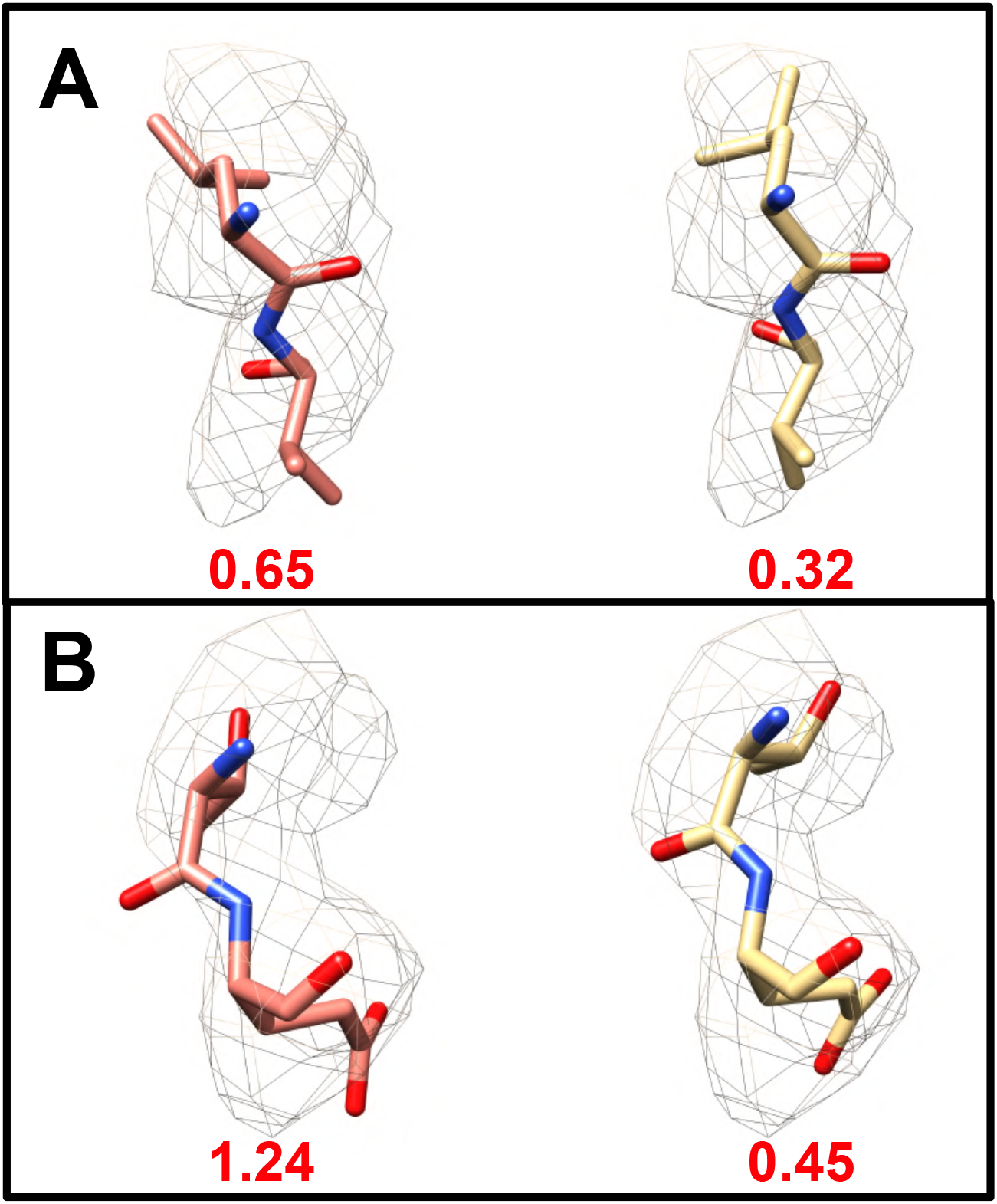
Improvement in the fit of the structure of the spike glycoprotein of SARS-CoV-2 in the closed state (emd-21452). In each panel, the left side shows the original fit and the right side shows the improved fit using ISOLDE (Croll, 2018). A) Fragment composed of amino acids LEU-533 and VAL-534 and B) THR-323 and GLU-324. Under each fragment the average FSC-Q is shown.

### 3.2. Overfitting

One of the big issues during the refinement of an atomic model is to detect and avoid overfitting. In the context of this work, overfitting occurs when the generated model is not supported by the reproducible signal on the half-maps. Different strategies have been presented to detect it. For example, the calculation of the FSC between one half-map and a map generated from the model refined against another half-map (DiMaio et al., 2013) or a comparison between the FSC of the original data and the FSC obtained using data with noise introduced at high resolutions (Chen et al., 2013). However, in both cases the problem is addressed globally.

The method we present here allows detecting the overfitting locally. Figure 1E and Figure 3A show that when the side chains are adjusted to densities corresponding to refinement artifacts, the FSC-Q values largely diverges from 0. Another example is shown in Figure 4, where the FSC-Q for PAC1 GPCR Receptor complex (emd-20278) (Liang et al., 2020) has been represented. Figure 4A shows several areas in red, which coincide with side chains that fall on densities corresponding to the detergent micelle. Examples of some of these amino acids are shown in the Figure 4B. In all these cases we observe that when the atoms fit to the detergent densities, FSC-Q moves away from zero and, importantly, FSC-Q gets negative values.

**Figure 3.**
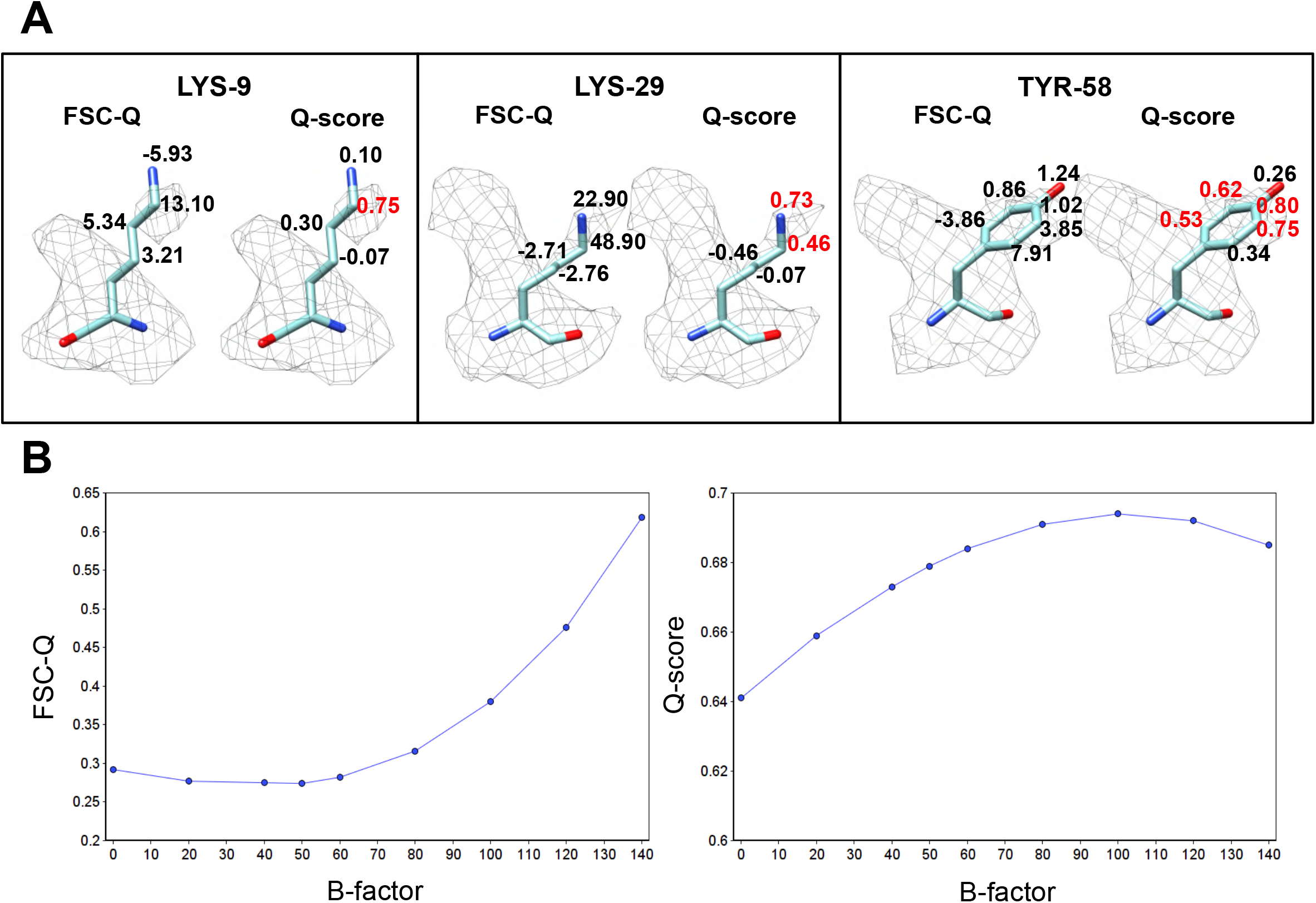
Comparison between FSC-Q and Q-score. A) FSC-Q and Q-scores for each side chain atom for the same residues as in Figure 1C. B) The plots show the effect of B-factor sharpening on FSC-Q and Q-scores analyzed using the 20S proteasome map.

**Figure 4.**
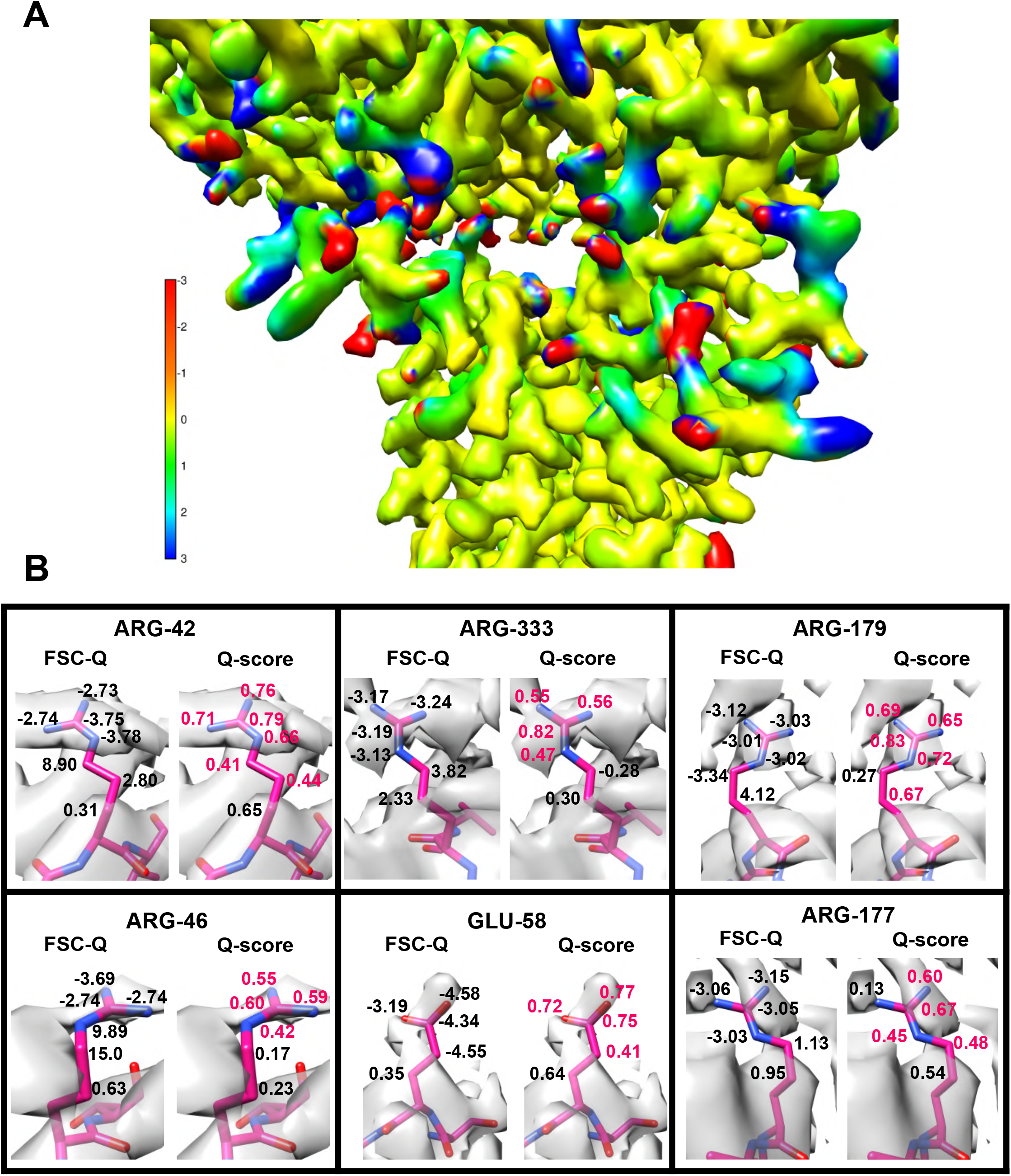
FSC-Q for a fragment of the PAC1 GPCR Receptor complex (emd-20278). A) FSC-Q values superimposed on the map generated from the atomic model (pdb id: 6p9y). Red areas indicate side chains that fit into densities of the detergent micelle. Blue areas correspond to atoms with very low resolvability. B) FSC-Q and Q-scores values are shown on each side chain atom of some residues that fit to the detergent [ARG-42 chain B, ARG-333 chain A, ARG-179 chain R, ARG-46 chain B, GLU-58 chain G and ARG-177 chain R].

### 3.3. FSC-Q as a complement to Q-score

The Q-score method measures the correlation between the map values at points around the atom and a reference Gaussian-like function (Pintilie et al., 2020). In this way, the Q-score estimates the resolvability of each atom with respect to the map from which the model has been refined. On the other hand, the method we present in this work measures how much of the model is supported by the signal in the half-maps. Additionally, we also include the environment of that atom within the local resolution window, spanning 5-7 times the global resolution. Although these methods measure different quantities, they both give a measure of the local quality-of-fit and complement each other.

The Q-score method is a good tool to estimate the resolvability of atoms. However, due to its normalization with respect to the mean, it is not capable of differentiating between the signal corresponding either to atoms or to densities generated by noise or artifacts of the reconstruction. Figure 3A shows the modified rotamers of the third block of Figure 1E with the FSC-Q and Q-score values for the atoms of the side chain. Several atoms are observed where the Q-score indicates a very good resolvability (in red color), however, these atoms are fitting into densities corresponding to noise. In contrast, the FSC-Q values for these atoms are very high, indicating a poor fit. This issue may have significant implications depending on the type of specimens. Particularly, for the very important biomedical case of membrane proteins, while FSC-Q is capable to identify the side chains where the fit is on the densities correspond to the detergent micelle (Figure 4), the Q-score is not, reporting values that would still show these fits as good (Figure 4B).

Both the Q-score method and our method depend on the B-factor that has been applied to the reconstructed map. In our method, this is due to the mask used in calculating the local resolution. For simplicity, we restrict ourselves to those cases in which sharpening is done globally, rather than on more precise local methods (such as Ramírez-Aportela et al. (Ramirez-Aportela et al., 2020), that, however, would complicate the presentation of these results significantly. To evaluate this dependence, the proteasome (emd-6287) was sharpened using different B-factor values, in the same way as in Ramírez-Aportela et al., 2019. In that paper we showed that a B-factor greater than −60 produced over-sharpening. The results obtained for the average value of Q-score and FSC-Q for each B-factor are graphically represented in Figure 3B. The charts show that FSC-Q decreases slightly, reaching the minimum approximately for a B-factor of −50 (in this particular case), with a difference of 0.018 with respect to the unsharpened map. From that point on, it begins to grow abruptly with the increase in the B-factor. This result indicates that as over-sharpening increases, FSC-Q increases. On the other hand, the charts show an increase in the Q-score, which reaches the maximum for a B-factor of −100, with a difference of 0.053 with respect to the unsharpened map. From there the Q-score decreases slightly. This indicates that the Q-score may correspond to some extent to oversharpening, and this effect may be a consequence of the normalization used.

## 4- Discussion

The ultimate goal of a cryoEM experiment is to generate an atomic model from the refined density map. The resolution of cryoEM maps may change in different areas and building an accurate atomic model using the map can be a real challenge. The development of metrics for map-to-model fit validation is a critical step towards robust generating high accuracy models from cryoEM maps.

In this paper, we proposed a new approach to analyze the fit of the models to the density maps. Our method is a significant extension of the map-to-model FSC calculation that is commonly used to assess goodness-of-fit (Lyumkis, 2019), since it explicitly introduces locality. Here we calculate the map-to-model local FSC and compare it with the half-maps local FSC. Differences between them allow us to select an objective signal threshold so that we can identify those areas where the model disagrees with the consistent signal on the half maps.

We have shown that FSC-Q enables identification of errors in fit and local detection of overfitting. Overfitting is one of the main challenges during model refinement and new metrics to detect it are needed (Lawson et al., 2020). The Q-score has been indicated by the authors (Pintilie et al., 2020) that is particularly sensitive to overfitting. Furthermore, we have shown that our method has less dependence on sharpening than the map Q-score and that they complement each other to better estimate the resolvability of atoms. Indeed, the combination of these (and other) methods probably will lead to meta validation measurements where the combination of methods as “orthogonal” as possible will lead to new simple metrics, yet to be explored, which could be directly incorporated into modeling workflows.

## Acknowledgments

We thank Prof. David Veesler for providing us the half maps of the spike glycoprotein of SARS-CoV-2. The authors would like to acknowledge financial support from: the Comunidad de Madrid through grant CAM (S2017/BMD-3817), the Spanish Ministry of Economy and Competitiveness through grant BIO2016-76400-R (AEI/FEDER, UE), the Instituto de Salud Carlos III, PT17/0009/0010 (ISCIII-SGEFI / ERDF) and the European Union and Horizon 2020 through grants: CORBEL (INFRADEV-01-2014-1, Proposal 654248), INSTRUCT-ULTRA (INFRADEV-03-016-2017, Proposal 731005), EOSC Life (INFRAEOSC-04-2018, Proposal: 824087) and HighResCells (ERC-2018-SyG, Proposal: 810057). The authors acknowledge the support and the use of resources of Instruct, a Landmark ESFRI project”. This work was supported by the Intramural Research Program of the National Institute for Arthritis, Musculoskeletal and Skin Diseases, NIH.

---

**Supplementary Figure 1.**
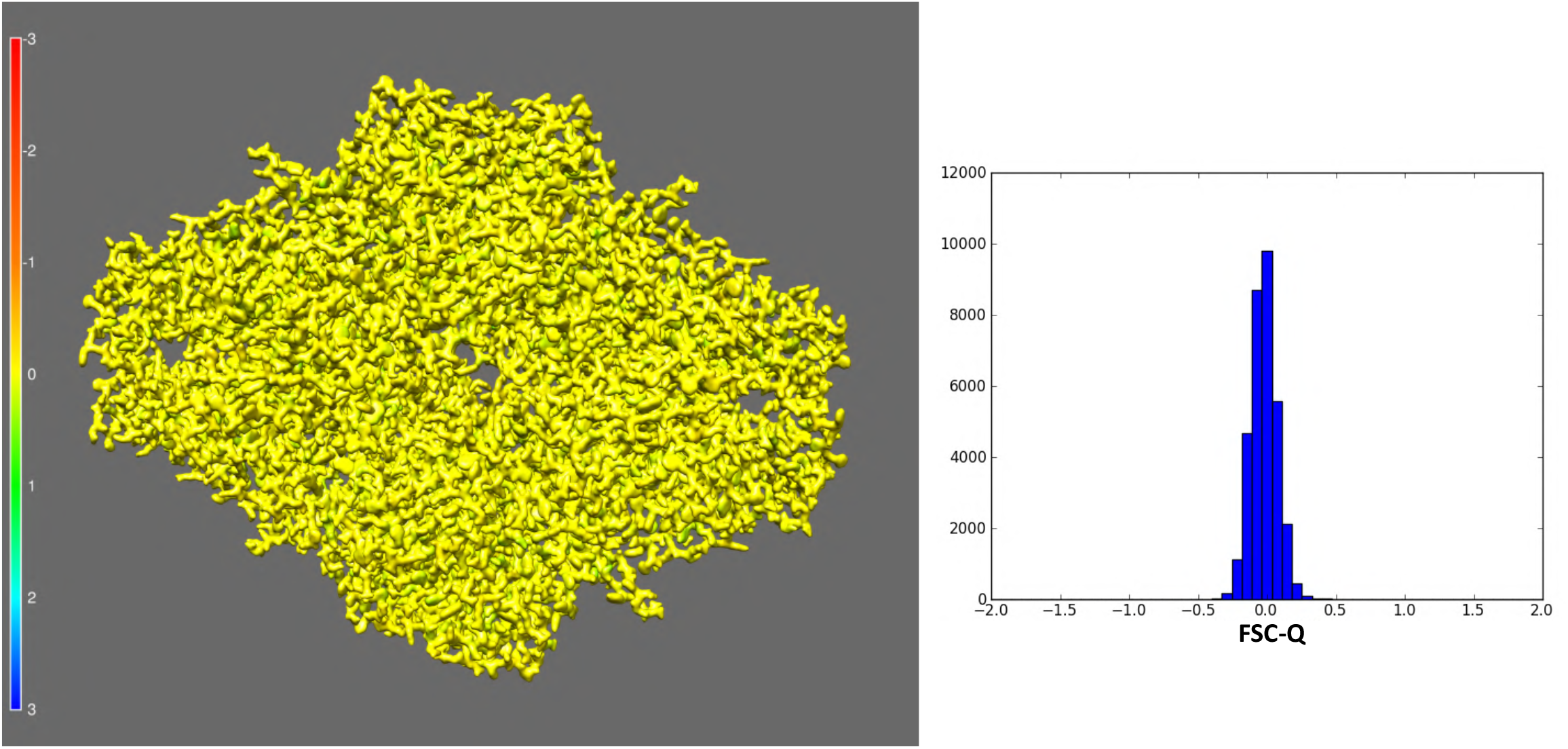
FSC-Q distribution for optimal map-to-model fit. For optimal fit, a map was reconstructed from the ß-galactosidase atomic model (pdb id: 3j7h), carrying out the following protocol on *Scipion*. First, a map was generated from the atomic model. Using the map, projections were generated in all directions with an angular sampling of 1.5 degrees, for a total of 18,309 projections. Gaussian noise with zero mean and a standar desviation of 50 was added to the set of projections and map reconstruction was carried out using the *RELION* software (Scheres, 2012).

